# Not all Bacterial Outer Membrane Proteins are β-Barrels

**DOI:** 10.1101/2023.10.25.564060

**Authors:** John Heido, Simon Keng, Pream Kadevari, Hulya Poyrazoglu, Ekta Priyam, Sajith Jayasinghe

## Abstract

The discovery of WzA, an octomeric helical barrel integral bacterial outer membrane protein, has challenged the widely held understanding that all integral outer membrane proteins of gram-negative bacteria are closed β-barrels composed of transmembrane β-strands. WzA is a member of the outer membrane polysaccharide exporter family and a bioinformatics analysis suggests that other members of the family may also contain outer membrane transmembrane segments that are helical. A review of the literature indicates that in addition to Wza, outer membrane core complex proteins of the type IV secretion systems also contain transmembrane segments that are helical.

## Introduction

Almost all integral membrane proteins found in the outer membrane of gram negative bacteria adopt a β-barrel architecture ^1,2^. These outer membrane proteins (OMPs) contain anti-parellel transmembrane (TM) beta-strands that are amphipathic with the non-polar face interacting with the hydrophobic region of the lipid membrane. Typically the TM b-strands of OMPs, as exemplified by OmpA ^3^, are segments of a single polypeptide chain with the N- and C-terminal stands interacting to form a closed barrel (Figure 1A). There is also an emerging class of β-barrel OMPs, represented by the Curli secretion channel CsgG ^4^, where the β-barrel is formed from TM β-strands contributed from individual subunits of a homo multimer (Figure 1B).

**Figure 1.**
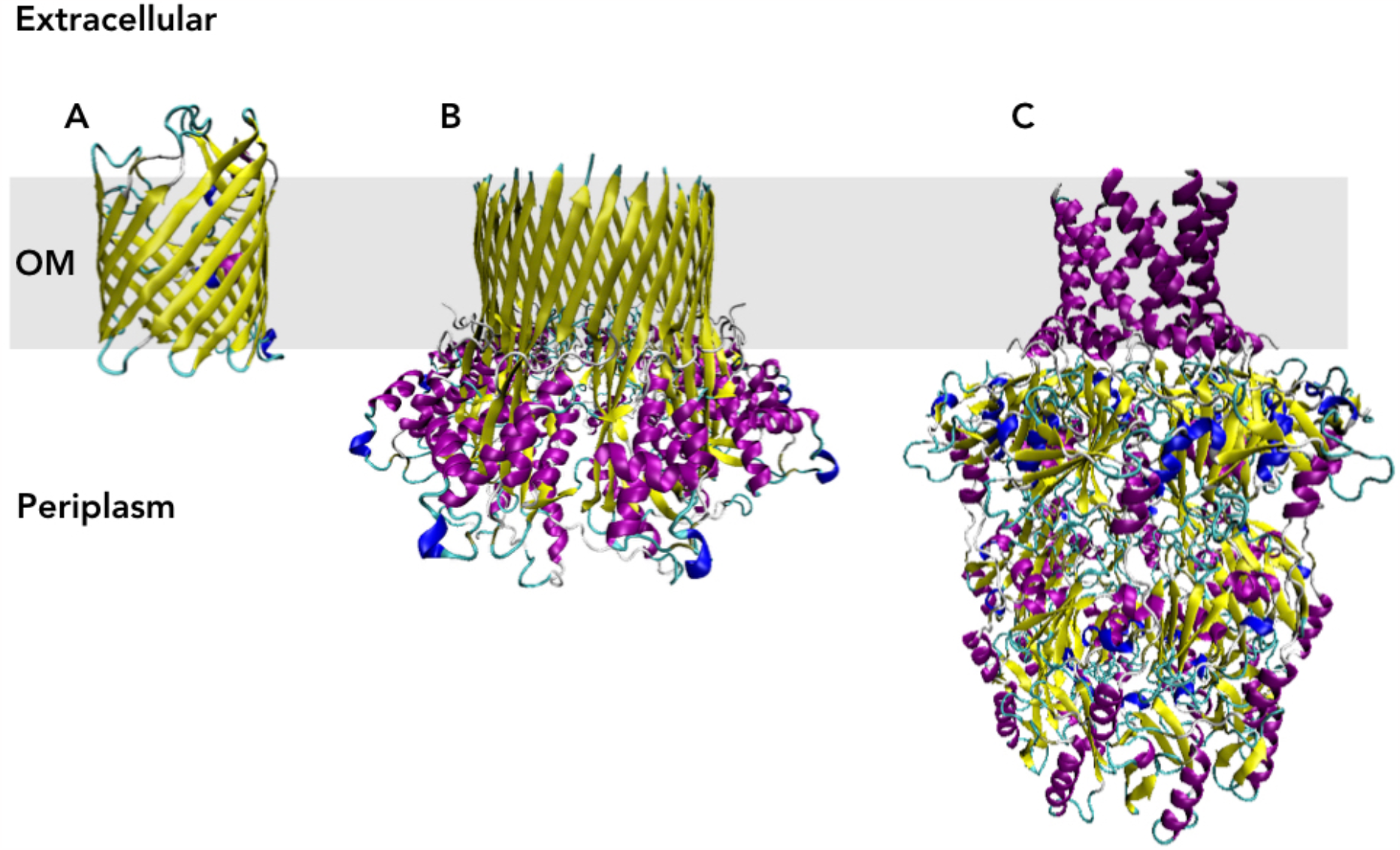
Representative structures of three classes of bacterial outer membrane proteins. (**A**). Structure of the porin OmpF (PDB ID: 2OMF monomer) where the transmembrane (TM) regions is composed of a 16-stranded β-barrel formed from a single polypeptide chain. (**B**). Structure of the curli secretion channel CsgG (PDB ID: 4UV3) where the TM region is composed of TM β-strands from nine individual polypeptide chains combining to form a 36-stranded β-barrel. (**C**). The structure of the group 1 capsular polysaccharide transporter WzA (PDB ID: 2J58) where the TM region is composed of α-helical TM segments from eight polypeptide chains combining to form a α-helical barrel.

It has long been thought that the outer membrane of gram-negative bacteria contains only β-barrel integral membrane proteins. This concept changed in 2006 with the discovery of Wza, an integral OMP involved in the transport of capsular polysaccharides ^5^. Based on the crystal structure, Dong et. al. proposed that Wza contains an α-helical c-terminal transmembrane segment. In the functional assembly the helical TM segments of eight Wza monomers associate to form an α-helical barrel structure that forms a pore in the outer membrane.

The discovery of WzA suggests that helical TM segments may be a more prevalent feature of gram-negative bacterial OMPs. A review of the literature and a bioinformatics analysis was carried out to identify other OMPs with helical TM segments.

## Materials and Methods

### Selection of Membrane Proteins and TM segments

Fifteen bitopic (single-spanning) bacterial inner membrane proteins were identified using the Membranome 3.0 database ^6^ and the TM segments of these proteins were identified in consultation with published accounts.

### Prediction of Helical Segments with Possible TM Segments

MPEx ^7^ and PsiPred (MEMSAT SVM) ^8,9^ were used to predict the presence of transmembrane segments. In the case of MPEx, predictions were carried out using the Wimley-White octanol scale ^10^ with a window size of 19 residues (defaults). PsiPred was used to predict the secondary structure distribution of each protein from which the helical secondary structure regions were identified.

### Determining Hydrophobicity and Amphipathicity of TM segments

Hydrophobicity, as qualified by the membrane insertion potential, ΔG_app_, of each TM segment was calculated using the ΔG prediction server at https://dgpred.cbr.su.se/index.php?p=home) ^11^. Amphipathicity, the asymmetric distribution of non-polar and polar amino acids, was quantified using the hydrophobic moment which was calculated using the translocon scale in the totalizer module of MPEx ^7^.

## Results and Discussion

### A c-terminal helical outer membrane TM segment may be common in OPX proteins

Integral membrane proteins of the bacterial outer membrane are typically β-barrel proteins containing β-strand TM segments. Until recently the one exception to this observation was the outer membrane protein Wza which contains a c-terminal helical TM segment. Wza belongs to the Outer membrane Polysaccharide Exporter (OPX), also known as the Outer Membrane Auxiliary (OMA), family of proteins { Cuthbertson:2009ip, Sande.2019 }. The OPX family of proteins are the final components in the transport of capsular and extra-cellular polysaccharides across the bacterial outer membrane, and are further subdivided into six groups (A through F) { Cuthbertson:2009ip }. Wza is a group A OPX and is involved in the transport of group 1 capsular polysaccharides. Although not all OPX protein appear to contain an outer membrane spanning domain ^14^ it is possible that those that do are anchored to the outer membrane via a helical TM segment. A sequence analysis was undertaken to determine if other OPX proteins might contain a c-terminal helix capable of anchoring these proteins to the bacterial outer membrane.

Thirteen protein sequences representing each of the OPX groups were compared to identify similarities that would support the hypothesis that other OPX proteins may have a structural organization similar to Wza. Pairwise sequence alignments of the thirteen OPX proteins indicates similarities ranging from 9 to 85 % between the different members of the family (Table 1). High sequence similarities (ranging from 40 to 85%) are observed for proteins within groups A, B, and C. Additionally, almost all members from OPX groups A, B, C, and D exhibit more than 25 % sequence similarity to Wza. Proteins belonging to groups E and F, which are much larger than the other OPX proteins, show less (∼ 18-22%) sequence similarity to Wza. The sequence similarly between Wza and the OPX proteins from groups A, B, C, and D suggest the possibility that proteins from these four groups could have a membrane architecture similar to that of Wza. This possibility is supported by the observation that all except two proteins (those from group D) have very similar secondary structure distributions to Wza (Figure 2 and also { Cuthbertson:2009ip }). The outer membrane helical TM segment of Wza is found at the extreme c-terminal region spanning residues 345 - 376 { Dong:2006el }. All of the group A, B, C, E and F proteins investigated here, but not the two from group D, contain a c-terminal helical region which could be embedded in the OM (Figure 2). Analysis of the protein sequences using MPEx { Snider:2009fl } and PsiPred ^9^ identified putative c-terminal TM regions, in all of the proteins investigated except the two representatives from group D (Table 2). These predicted TM segments correspond to the c-terminal helical regions identified using secondary structure analysis. Taken together these observations point to the distinct possibility that at least group A, B, C, E and F OPX family members may also have an a-helical transmembrane segment which helps to form a pore in the bacterial outer membrane.

**Table 1.**
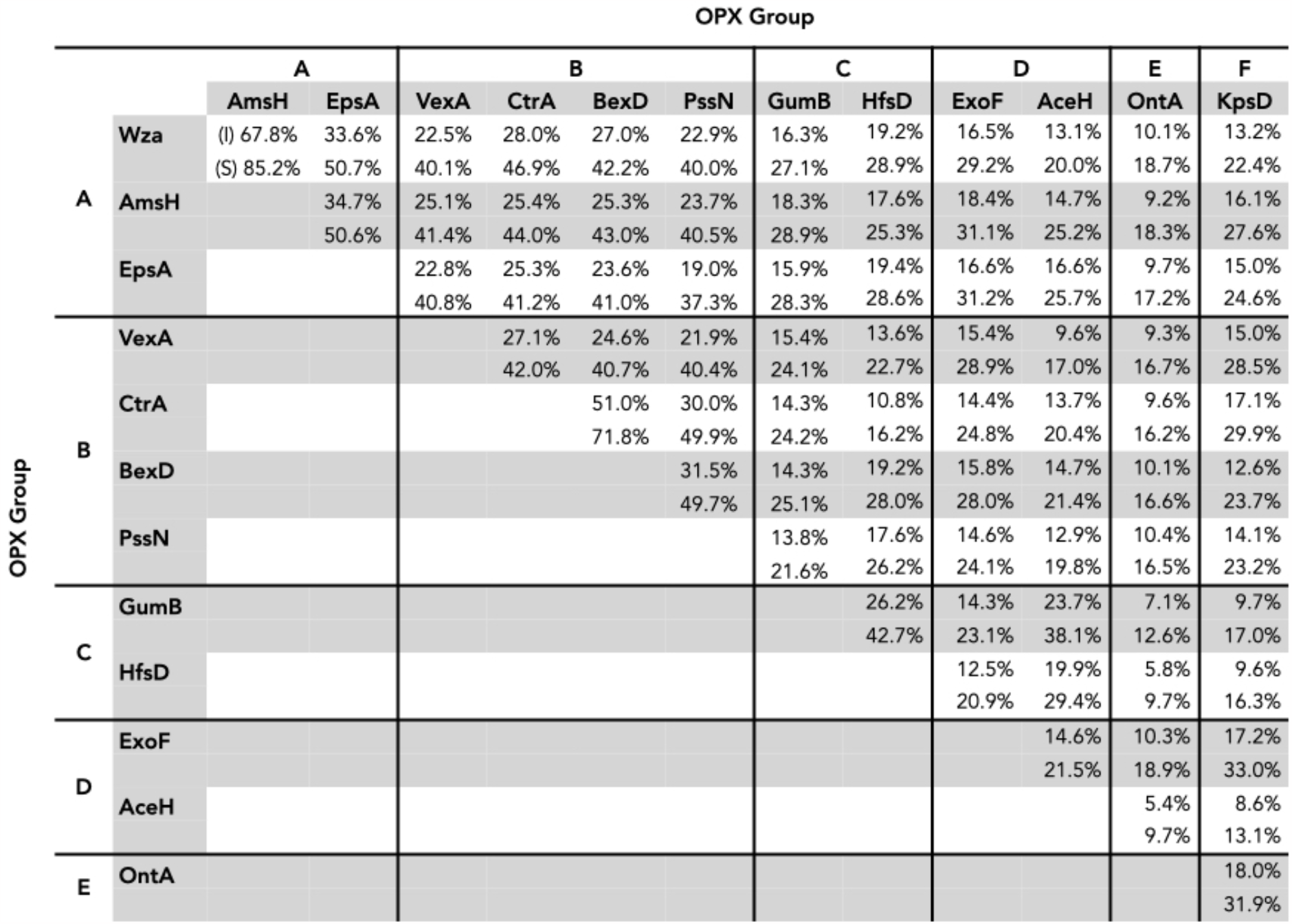
Sequence identity (I) and similarity (S) of group A, B, C, D, F, and F OPX proteins. Group A: Wza (UniProt-P0A930), AmsH (Q46629), EpsA (Q45407); Group B: BexD (P22236), CtrA (P0A0V9), PssN (Q27SU9), VexA Q04976); Group C: GumB (Q456768), HfsD (Q(A5L5); Group D: ExoF (Q02728), AceH (Q8RR80); Group E: OntA (Q56653); Group F: KpsD (Q03961). OPX proteins from groups A, B, C, and D show high sequence similarity to each other (ranging from 40 to 85%) and exhibit more than 25 % sequence similarity to Wza. Group E and F proteins are much larger than other OPX proteins and exhibit less similarity to Wza.

**Tables 2.**
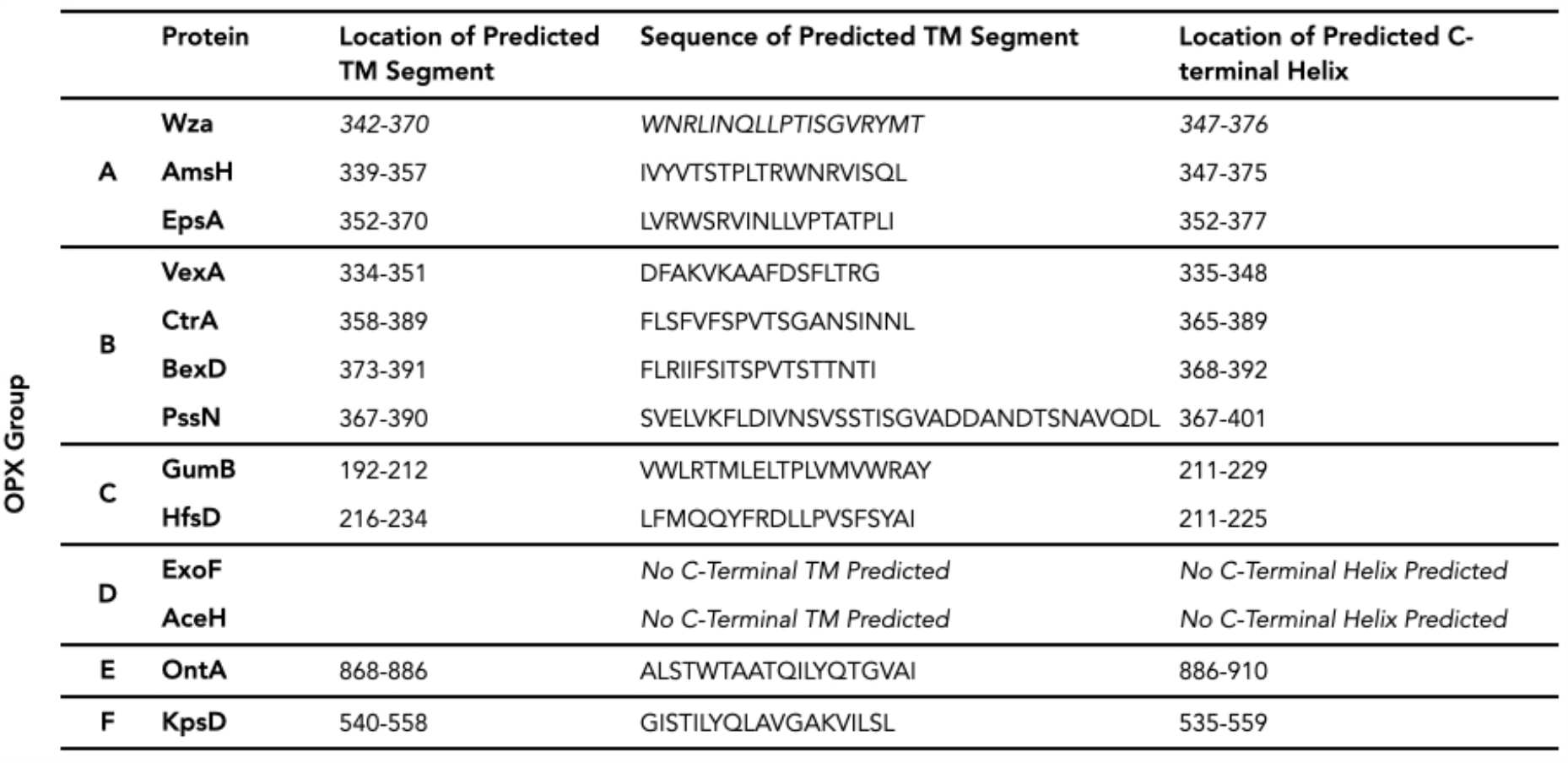
C-terminal helical TM segments predicted in representative examples of OPX proteins. The transmembrane segment of Wza, the only OPX protein whose structure is known, is located at the c-terminus of the protein. We used PsiPred (MEMSAT SVM) ^9^ and MPEx { Snider:2009fl} to identify any possible c-terminal TM segments present in OPX proteins. We also predicted the secondary structure distribution of these proteins using PsiPred and identified the most c-termial helix. All of the proteins investigated, except the two from group D, contain a c-terminal helical region and these are predicted to be possible TM regions.

**Figure 2.**
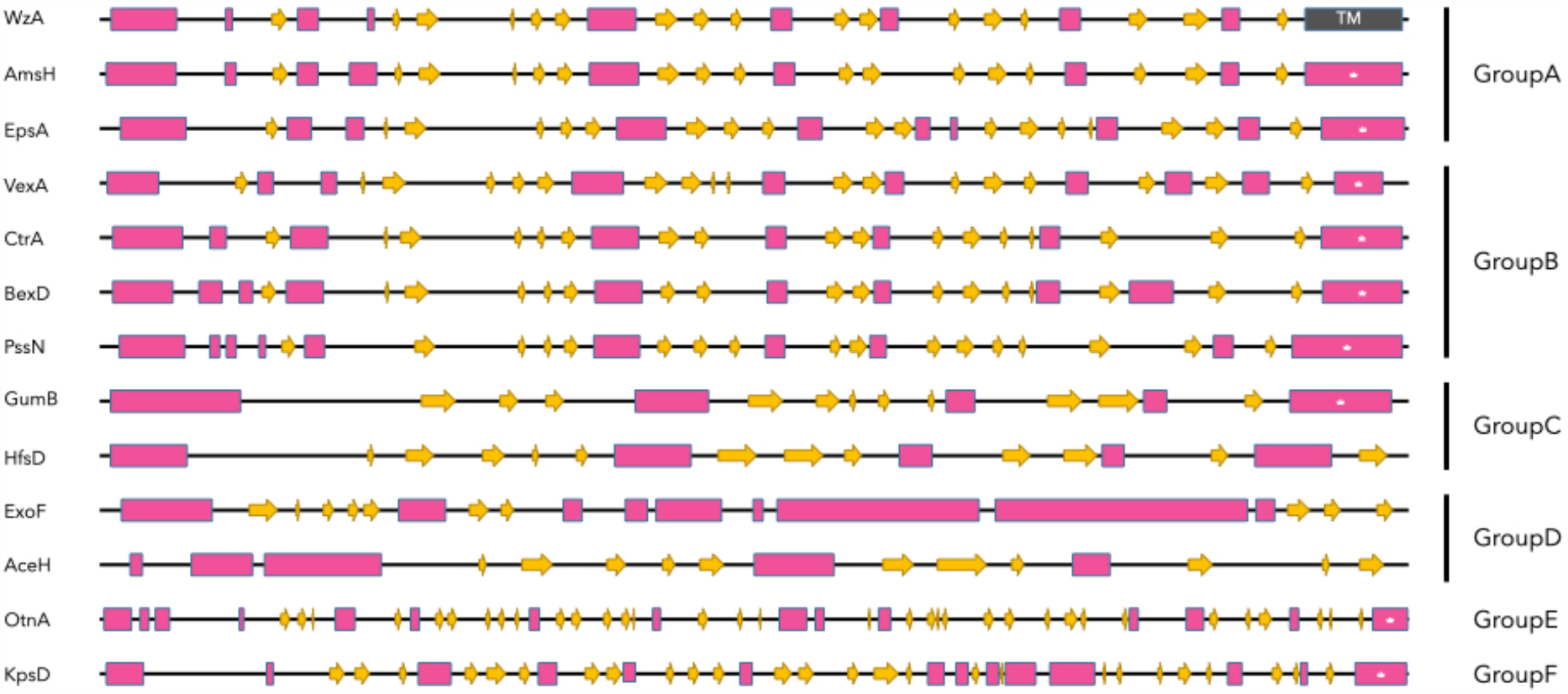
Predicted secondary structure distribution of thirteen OPX proteins representing the six OPX classes. β-strand are shown as yellow arrows while helices are shown as pink rectangles. Secondary structure distribution was predicted using PsiPred ^9^. The helical TM segment of Wza, the only OPX protein whose structure is known, is indicated as a grey rectangle. All proteins expect ExoF and AceH from group D indicate the presence of a c-terminal segment (*). MPEx and PsiPred also predict these regions to be possible TM segments.

### A helical TM segment is also found in the outer membrane pore component of type IV section systems

In addition to Wza, at least six other proteins, all outer membrane core complex components of the type IV secretion system (T4SS), have been identified as containing helical TM segments. T4SSs are a diverse family of bacterial protein complexes that mediate the transfer of DNA and protein components to target cells ^16–19^. The canonical T4SS complex from *Agrobacterium tumefaciens* is composed of 12 proteins, named VirB1 to 11 and VirD4, that spans the innermembrane, the periplasm and the outer membrane. The outer membrane core is composed of three proteins, VirB7, VirB9, and VirB10, of which VirB10 forms an outer membrane pore { Sgro:2019hb }. In the case of VirB10 from *Xanthomonas citri* { Sgro:2018bh }, the outer membrane pore is composed of two concentric rings of c-terminal TM helices from 14 VirB10 molecules, where the α-helices of the outer-ring contact the membrane while the helices of the inner ring line the pore (Figure 3A). A similar architecture of TM helices is observed in VirB10 equivalents from the outer membrane core complexes of at least five other type IV secretion systems: *E. coli* TraF, a VirB10 homolog encoded by the pKM101 plasmid { Chandran:2009fg }, *E. coli* TrwE, a VirB10 homolog encoded by the R388 plasmid { Macé.2022 }, *Salmonella* TraB, a VirB10 homolog encoded by the F plasmid { Liu.2022, Amin.2021 }, CagY, the VirB10 homolog from *Helicobacter pylori* { Chung:2019fz }, and DotG, the VirB10 homolog from *Legionella pneumpphila* { Sheedlo.2021 }. Despite the presence of a wide variety of T4SSs, VirB10 appears to be a core component of the assembly, and the C-terminus of these proteins appears to be conserved { Sheedlo.2021 }. Thus the presence of c-terminal helical TM segments may be a common feature of T4SSs.

**Figure 3.**
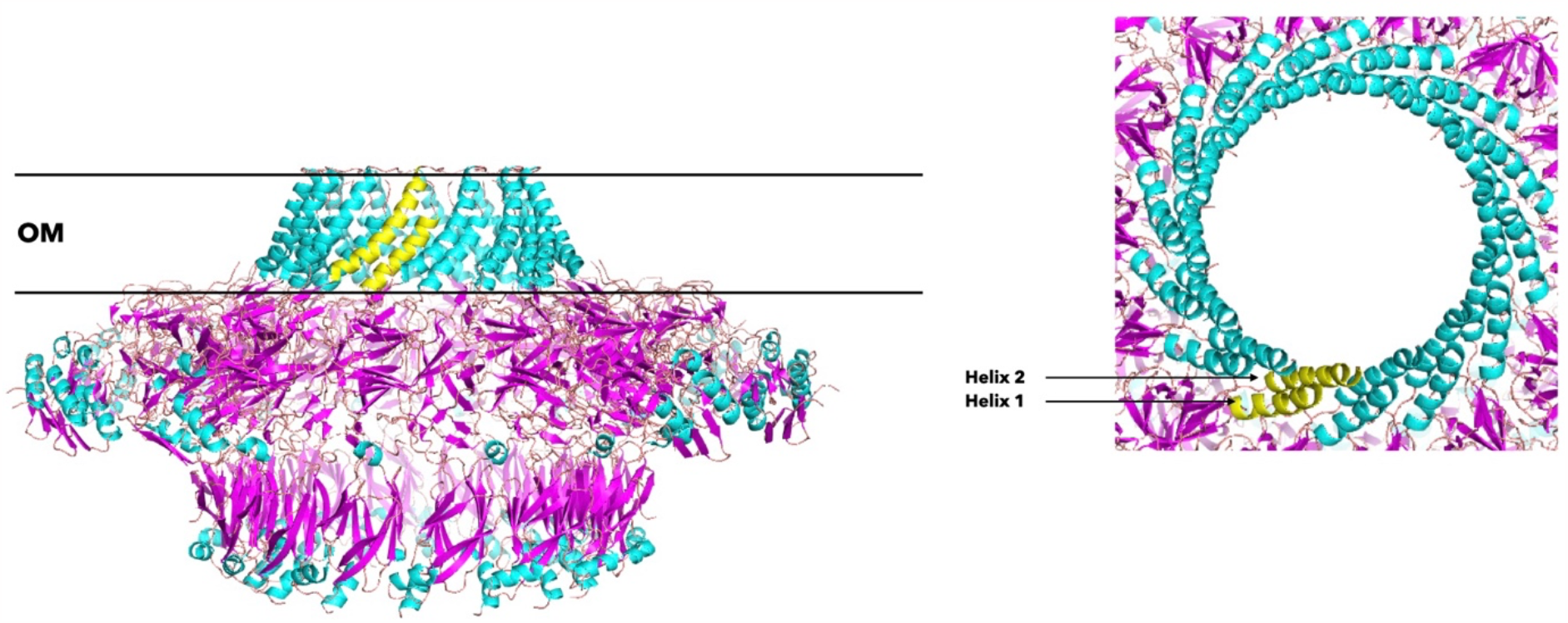
Structure of the outer membrane core complex of T4SS of *Xanthomonas citri (*PDB ID 6GYB) { Sgro:2018bh}. Side (left) and top (right) view of the complex composed of VirB9 and VirB10 are shown in ribbon representation. The outer membrane α-helical transmembrane segments of VirB10 are shown in yellow. The TM helices form two concentric rings that line the pore of the T4SS outer membrane channel. VirB10 appears to be present in nearly all T4SS complexes { Sheedlo.2022} and therefore, the presence of helical outer membrane segments may be a common feature of T4SSs.

### Helical TM segments of outer membrane proteins are in general less hydrophobic and more amphipathic than helical TM segments of inner-membrane proteins

Membrane proteins destined for the bacterial outer membrane must first pass through the inner membrane. One route through the inner membrane involves the SecYEG translocon which helps the transport of proteins and is also responsible for facilitating the insertion of proteins into the inner membrane. The insertion of TM segments into the inner membrane by SecYEG appears to involve partitioning of protein segments between the translocon channel and the lipid membrane, and it is thought hydrophobicity plays a role in the identification of segments destined to become inner membrane TMs ^27,28^. If the helical TM segments of outer membrane proteins traverse through the SecYEG translocon their hydrophobicity must be sufficiently low to avoid accidental insertion into the IM.

The potential for a given amino acid sequence to be recognized as a TM and be inserted into the inner membrane, the membrane insertion potential (ΔG_app_) can be estimated ^11^. Based on the observation that while TM segments of biotopic (single spanning) integral membrane proteins almost always exhibit ΔG_app_< 0 kcal/mol {Hessa 2007} and secreted proteins almost never contain segments with ΔG_app_< 0 { Hessa:2007dk }, it is generally taken that an amino acid segment with a negative ΔG_app_ will be recognized by the translocon to be TM and be inserted in to the inner membrane. We compared ΔG_app_ values of the alpha-helical TM segments of outer membrane OPX and Type IV secretion system proteins to values from helical TM segments of a collection of biotopic integral inner membrane proteins. All of the helical TM segments from the outer membrane proteins exhibit ΔG_app_>0, whereas almost all of the TM segments from bitopic bacterial inner membrane proteins exhibit ΔG_app_< 0, suggesting that the helical TM segments from outer membrane proteins investigated here have low inner membrane insertion potential (Figure 4A). This observation supports our expectation that low membrane insertion potential, or hydrophobicity, allows outer membrane helical TM segments to pass through the translocon without accidentally being inserted into the inner membrane.

**Figure 4.**
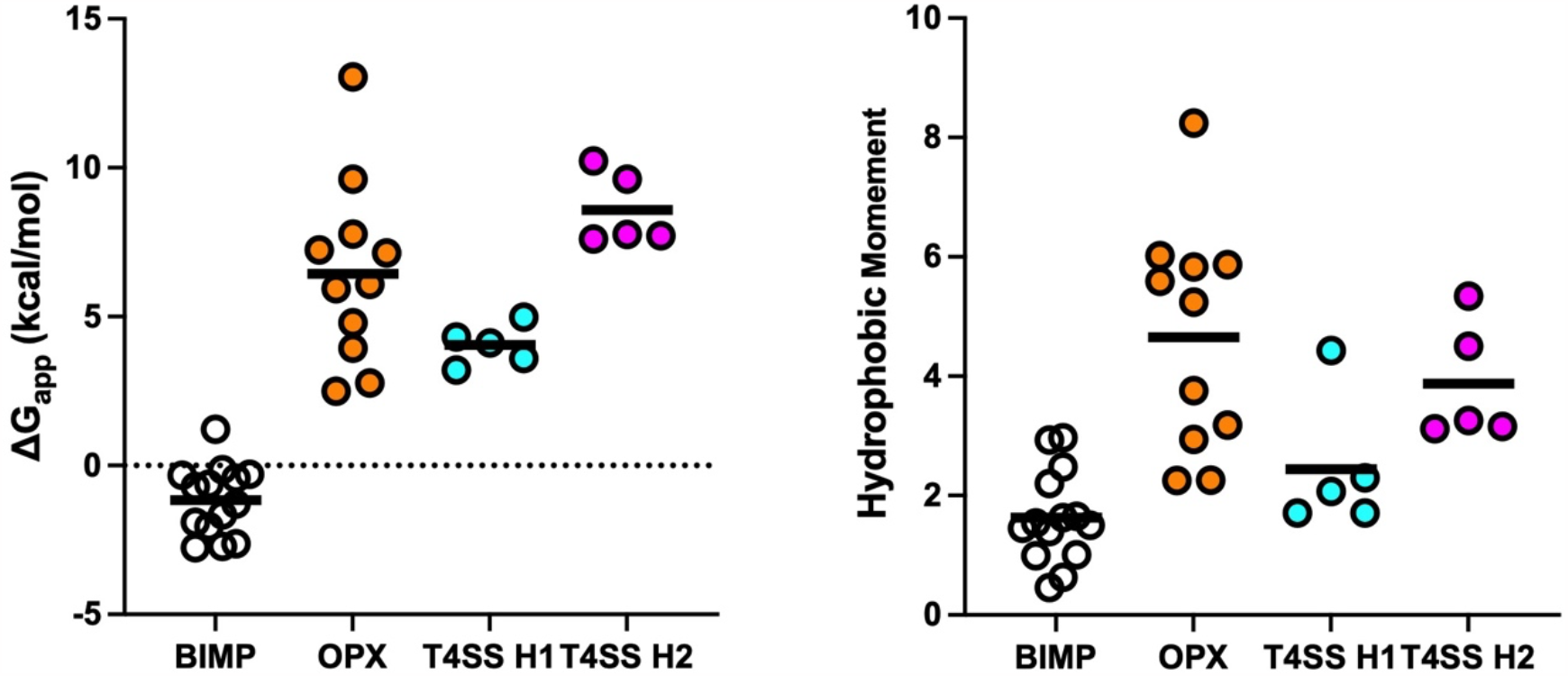
Inner membrane insertion potential (ΔG_app_) and hydrophobic moment (amphipathicity) of helical TM segments of outer membrane proteins. **(A)** Almost all of the TM segments biotic inner membrane proteins (BIMP) inserted into the bacterial inner membrane (by the translohcon) exhibit ΔG_app_ <0 (A, black circles, n=14, -1.12 ± 0.31) while the ΔG_app_ values of the α-helical TM segments of outer membrane OPX (A, orange circles, n=11, 6.44 ± 0.93) and T4SS proteins (A, blue for helix 1, n=5, 4.04 ± 0.30, and pink, for helix 2, circles, n=5, 8.59 ± 0.56) have ΔG_app_>0 suggesting that the helical TM segments from the outer membrane proteins investigated here have low inner membrane insertion potential. **(B)** helical TM segments from the OPX proteins (orange circles, n=11, 4.65 ± 0.57) and TM helix 2 (pink circles, n=5, 3.88 ± 0.45) of the T4SSs have hydrophobic moments that are on average larger than those observed for the helical TM segments of biopic inner membrane proteins (black circles, n=14, 1.63 ± 0.21). Outer membrane TM helix 1 (blue circles, n=5, 2.44 ± 0.51) of the T4SS system is much less amphipathic than TM helix 2. It has been suggested that the amphipathic nature of the helical TM segments facilitate outer membrane insertion ^37^.

It should be noted that a positive ΔG_app_ value does not by itself preclude insertion into the inner membrane. In fact a significant number of TM segments from polytopic integral membrane proteins have been found to have quite positive ΔG_app_ values, to the extent that some segments are not even initially recognized as TM by the translocon {Marothy 2015, Hedin 2010,Whitley 2021}. The insertion of TM segments with positive ΔG_app_ appears to depend on surrounding sequence features such as the presence of flanking charged amino acids and the presence of TM segments with high insertion potential { Marothy.2015, Hedin:2010he }. Such features are lacking in the outer membrane proteins discussed here presumably ensuring that the TM segments of these proteins pass through the translocon.

Once proteins destined for the outer membrane pass through the translocon they are transported through the periplasm and targeted to the bacterial outer membrane at which point the TM segments insert into the outer membrane. In the case of integral beta-barrel proteins, which comprise the majority of outer membrane proteins, the details of this process are established and integration into the outer membrane involves the BAM complex ^34^. Although mechanistic details of the integration of outer membrane proteins with helical TM segments is not known, at least in the case of Wza the process does not appear to involve the BAM complex { Dunstan:2015jw }. It has been proposed that integration of outer membrane proteins that do not contain a traditional beta-barrel architecture follows a novel pathway independent of BAM { Dunstan:2015jw } and that perhaps the amphipathic TM regions of these proteins facilitate membrane insertion ^37^. The TM segments of OPX and T4SS proteins are amphipathic (Figure 4B), especially those from the OPX proteins and TM helix 2 of the T4SSs, which have hydrophobic moments that are on average larger than those found in bitopic inner membrane protein TM segments. The helical TM segments of the OPX and T4SS proteins form membrane channels and we expect their hydrophilic and hydrophobic amino acids to separate to form membrane interacting and pore lining regions respectively resulting in a large hydrophobic moment.

Although the vast majority of proteins embedded in the bacterial outer membrane adopt a β-barrel architecture members of at least two classes of proteins appear to contain helical TM segments embedded in the outer membrane. The identification of these proteins changes the long held understanding that all bacterial outer membrane proteins were β-barrels and opens up the possibility that other classes of outer membrane proteins may also contain TM helices.

## Acknowledgements

We thank current laboratory members for editing the manuscript.

## Figure Legends

Figure 1. Representative structures of three classes of bacterial outer membrane proteins. (**A**). Structure of the porin OmpF (PDB ID: 2OMF monomer) where the transmembrane (TM) regions is composed of a 16-stranded β-barrel formed from a single polypeptide chain. (**B**). Structure of the curli secretion channel CsgG (PDB ID: 4UV3) where the TM region is composed of TM β-strands from nine individual polypeptide chains combining to form a 36-stranded β-barrel. (**C**). The structure of the group 1 capsular polysaccharide transporter WzA (PDB ID: 2J58) where the TM region is composed of α-helical TM segments from eight polypeptide chains combining to form a α-helical barrel.

Figure 2. Predicted secondary structure distribution of thirteen OPX proteins representing the six OPX classes. β-strands are shown as yellow arrows while helices are shown as pink rectangles. Secondary structure predicted using PsiPred ^9^. All proteins expect ExoF and AceH from group D indicate the presence of a c-terminal helical segment (*). MPEx and PsiPred also predict these regions to be possible TM segments.

Figure 3. Structure of the outer membrane core complex of T4SS of *Xanthomonas citri (*PDB ID 6GYB) { Sgro:2018bh}. Side (left) and top (right) view of the complex composed of VirB9 and VirB10 are shown in ribbon representation. The outer membrane α-helical transmembrane segments of VirB10 are shown in yellow. The TM helices form two concentric rings that line the pore of the T4SS outer membrane channel. VirB10 appears to be present in nearly all T4SS complexes { Sheedlo.2022} and therefore, the presence of helical outer membrane segments may be a common feature of T4SSs.

